# Assessing suitability of ultra-low-field MRI for TES electric field models

**DOI:** 10.64898/2026.01.06.697579

**Authors:** Alexander R Guillen, Yu Huang, Dennis Q Truong, Niranjan Khadka, Yishai Z Valter, Christopher C Abbott, Zhi-De Deng, Abhishek Datta

## Abstract

**Objective:** Recent ultra-low-field (ULF) MRI systems provide an option for more accessible and cost-effective imaging to widely used higher field (HF, usually 3T) systems. However, it remains uncertain whether ULF scans can be used for reliable TES current-flow modeling as opposed to using 3T MRI.

**Methods:** We used raw anatomical scans of 23 healthy adults from 64 mT and 3T systems. We enhanced the 64 mT scans using SynthSR. Using our established current-flow modeling pipeline we determined the electric fields (EF) for the ULF and 3T scans induced by the classic M1-SO electrode montage. We quantified the differences in EF at three regions of interest (ROI): left motor cortex (LMC), anterior commissure (AC), and left hippocampus (LHC).

**Results:** At the cortical ROI (LMC), the individual mean EF based on a 3T scan was found to be 27% higher than the corresponding ULF scan. The average mean relative error was found to be 40%. At AC, the individual mean EF based on a 3T scan continued to be elevated by about 13% in magnitude with respect to the corresponding ULF scan with a mean error range of 31-34%. At LHC, comparison of the induced individual mean EF magnitudes indicated no clear bias between 3T-based and ULF-based models. The average mean relative error at this level dropped to ∼20%.

**Discussion:** The substantial decrease in cortical EF when using an ULF scan is predominantly explained by the overestimation of cerebrospinal fluid (CSF) noted by prior morphological comparison studies. Scaling up EF magnitude to match predicted cortical values to a 3T scan may be considered depending on the modeling question being addressed. While preliminary, our results suggest an “equalizing” effect between ULF and 3T predictions with increasing depth within the brain.

## 1. Introduction

The recent advent of ultra-low-field (ULF) MRI scanning has the potential to transform neuroimaging. Powered by a standard electrical socket, portable form factor, and low cost, it can enable access across a wide range of settings (community and resource-limited) (1).

Transcranial electrical stimulation (TES) is a non-invasive modality that typically delivers low intensity current (∼2 mA) using two scalp electrodes to modulate the underlying neuronal excitability (2). This modulation of membrane excitability ultimately determines observed behavioral / clinical outcomes. Owing to current-controlled application, physics dictates that current entering from one electrode has to exit from the second electrode (3). In case of bipolar waves, current simply reverses between the two electrodes every cycle. Computational models using electric fields (EF) are a standard tool to simulate and visualize this current-flow propagation (4,5).

Such modelling enables researchers to understand which regions receive less or more “dose” as it is rational to assume that regions subject to higher current flow are more likely candidates for modulation and plasticity. Consequently modeling has been used to make stimulation / targeting decisions, retrospectively analyze stimulation outcomes (4), unravel dose-response relationships, and study safety implications (6). Individualized models are developed directly based on an individual’s structural MRI (7,8). Specifically, the MRI is segmented into different tissues based on gray scale values and subsequently meshed to create an anatomically valid head model. Therefore the accuracy of the computational modeling process is critically dependent on the quality of the MRI scan (5). Typical modeling workflows since ∼2009 have employed scans obtained from a higher field (HF) of over 1.5T (7,9). Further, modeling methodology based on 3T scans has been validated using intracranial recordings demonstrating its utility (10).

With the imminent wider use of ULF systems, it becomes naturally important to assess the suitability of these scans for TES modeling. Vasa et al recently reported on the morphometric correspondence of ULF (64 mT) to high-field MRI (3T) measurements using correlations of tissue volume (11). When comparing percentage differences, authors noted median underestimates in both gray matter (GM) and white matter (WM) (GM cortical= −1.5%, GM subcortical = −3.7%, WM= −4.4%) and overestimates in cerebrospinal fluid (CSF) (+24.6%). The authors noted high tissue volume correlations in larger structures (Pearson’s r:0.95-1) but with Lin’s concordance correlation coefficient (CCC) in the range 0.39-0.95. The correspondence at the individual structure level was also reported to be high but in general, larger structures were shown to have a higher correspondence to 3T. With respect to implication for TES modeling, it is well known that the individual CSF anatomy enclosing / perfusing the gyrated cortex plays a defining role in induced current-flow patterns. For instance, a particular gyrus may be subject to high induced EF due to: a) wide pockets of CSF on either side and/or b) reduced CSF thickness concentrating or “funneling” current into that gyri crown (7). On the contrary, a thicker layer of CSF over a particular gyrus will result in low induced EF in the gyrus due to motivation for current to shunt away (i.e. traverse across the higher conductive CSF). Given the high overestimation of CSF reported in ULF scans, it is therefore rational to expect an impact in the current-flow patterns with respect to the corresponding HF scans.

In this exploratory study, we sought to investigate and quantify differences in induced EF between subject-matched ULF- and 3T-based models using the finite element method (4,7). We considered the classic M1-SO electrode montage used in TES (12). We evaluated differences at three different regions of interest (ROI). Given the intended brain target for the M1-SO is the underlying cortical region under the M1 electrode, we considered the left motor cortex (LMC). We further evaluated differences at the anterior commissure (AC), a classic neuroimaging reference point and left hippocampus (LHC), a popular subcortical region. For all three ROIs, we considered spherical volumes of 5 and 10 mm radii. Our ultimate goal was to study differences and assess the impact of using ULF scans.

## 2. Materials and Methods

### 2.1 Dataset

The structural scans used in this study were obtained from the publicly posted dataset by Vasa and colleagues in the context of their ULF brain morphometry study (11). As opposed to prior work on correspondence of 64 mT morphometric measurements to 3T(13), the dataset from Vasa et al (11) is novel as it did not utilize image quality transfer, eliminating potential for artifacts introduced by neural networks. The defaced dataset comprised 23 healthy subjects (11 females, 12 males, age of 20 to 69 years) and were balanced for age and sex, with a minimum of 2 male and 2 female head scans available in each decade of age. We considered the ULF dataset (“ses-HFC”) that was acquired on the same day when the HF dataset was acquired. The 3T MRI data consisted of both T1-weighted (T1w) and T2-weighted (T2w) scans acquired at 1 mm isotropic resolution. The ULF data available was anisotropic with the following resolutions – T1w: 1.6×1.6×5 mm^3^; and T2w: 1.6×1.6×5 mm^3^.

### 2.2 Pre-processing

The ULF scans were corrected using SynthSR (14) using corresponding T1w and T2w images to enhance image quality. As the face was removed in both the ULF and 3T datasets, the AFNI refacing algorithm was applied to recreate facial features (15). Refacing was particularly important as we are studying the induced EF from the M1-SO electrode montage, where the supraorbital (SO) electrode was positioned on the upper face that was removed in the raw data. To avoid complications in electrode placement, residual artifacts in ULF scans near the head vertex were removed by assigning them a value of zero.

### 2.3 EF computation

Forward modeling of the EF was performed using ROAST (16) with a 5×5 cm electrode pad in the M1-SO configuration at 2 mA. Specifically, electrodes at the C3 and Fp2 locations in the international 10-10 EEG convention (17) were used when calling the roast() function, and the default pad orientation was used (long axis of the pad going left to right). Note that the anisotropic ULF data was converted into isotropic 1 mm resolution by SynthSR, and thus no resampling was performed when calling the roast() function. We used the default conductivity values for the modeled tissues and electrodes (in S/m): skin: 0.465; skull: 0.01; CSF: 1.65; gray matter: 0.276; white matter: 0.126; air: 2.5 × 10^-14^; electrode: 5.8 × 10^7^; gel: 0.3).

### 2.4 Post-processing

The segmentation of head tissues for each subject generated from ROAST was imported into Simpleware (Synopsys Ltd., CA, United States) for visualization in 2D and 3D (**Figure 1**). To observe the difference in tissue segmentation, we visualized the exemplary subjects spanning the extremes of the age from the dataset (Subj01 & Subj04: 20-29 years; Subj11 & Subj20: 60-69 years).

**Figure 1:**
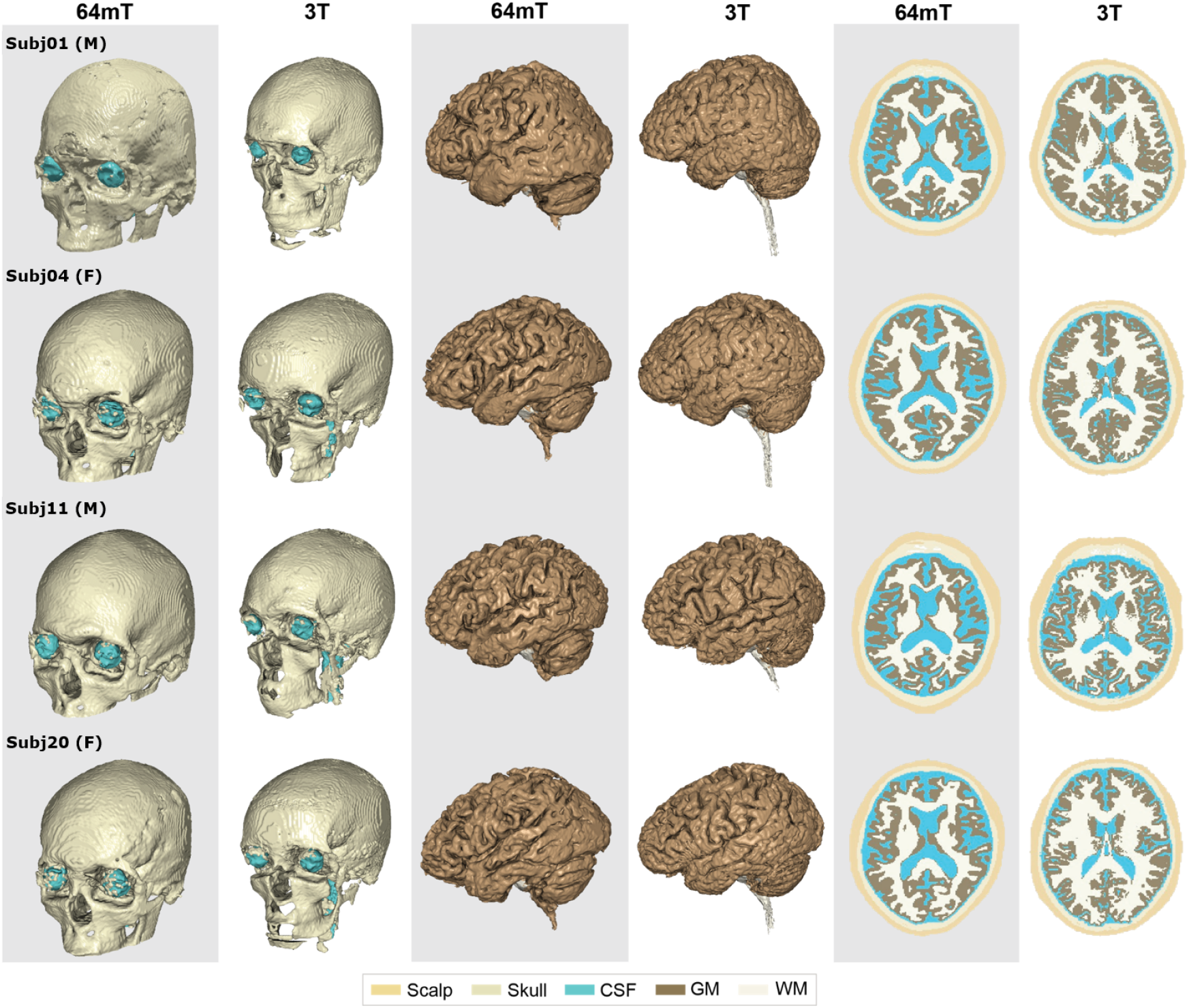
Illustration of segmentation masks rendered in 3D and 2D for four subjects spanning extremes of age (20-29 years and 60-69 years). The data corresponding to the ULF scans is shown in gray background. The first and second columns, and the third and fourth columns, represent segmented skull and gray matter masks from ULF (64mT) and 3T MRI scans, respectively. The fifth and sixth column represent an axial cross section (at the level of anterior commissure) of the segmented masks from ULF and 3T MRI scans, respectively. Depicted segmentated tissue masks are shown using colored blocks at the bottom.

Post-processing of the EF was conducted in MATLAB (MathWorks, MA, United States). Specifically, we are interested in how the EFs computed from ULF scans are different from those in the 3T scans, at three ROIs: LMC (MNI coordinates (−50, −14, 46)), AC (0, 0, 0), and LHC (−26, −12, −14) (18). As ULF and 3T scans have different sizes in their native space, and warping the computed EF from ROAST to a common space may distort the field vectors, we compared the EF across the two scanning modalities at ROIs mapped from the MNI space. Specifically, a sphere of radius of either 5 mm or 10 mm at one of the ROIs in the MNI space was mapped to the native space of both the ULF and 3T scans, using an affine transform estimated by the Unified Segmentation algorithm (19) implemented in the SPM software package that was called by ROAST. The mapped sphere was checked against the brain mask output by ROAST, and locations in the mapped sphere that are outside of the brain were discarded. EF magnitude and vectors were read out at the sphere, and the mean and standard deviation of the field magnitude in the sphere were calculated, for each modality and each subject. To quantify the vectorial difference of the EFs from the two modalities, we also computed the relative difference (rD) of the EFs as: rD = |**E**_**ulf**_ - **E**_**3T**_| / |**E**_**3T**_|, where **E**_**ulf**_ and **E**_**3T**_ are the field vectors from ULF and 3T scans, respectively, and || is the vector norm. The relative difference was calculated voxel by voxel at the locations shared by the two modalities.

## 3. Results

### 3.1 Differences in tissue segmentation between the two imaging modalities

**Figure 1** illustrates tissue masks from exemplary subjects spanning the extremes of the age from the dataset evaluated. We observed no striking differences visually with respect to scalp (not shown) and skull discontinuities. We further observed no discontinuities in the CSF layer indicating that our segmentation pipeline is able to appropriately perform tissue assignment on processed ULF scans. We however note clear CSF overestimation for the ULF scan in comparison to its corresponding 3T scan. We further observe subtle overestimation of the GM as well. Overall, individual tissue anatomy / details are preserved including zygomatic arch/process, foramen magnum, gyri folding, etc.)

### 3.2 Differences in the induced electric field (EF)

The *individual* mean EF in the LMC for a model based on 3T scan was found to be higher than the corresponding ULF scan, for both sizes of ROI (5 mm, ULF vs 3T: 0.22 ± 0.04 V/m vs 0.28 ± 0.06 V/m, **Figure 2A**; 10 mm, ULF vs 3T: 0.22 ± 0.04 V/m vs 0.29 ± 0.06 V/m, **Figure 2B**). This indicates that forward models based on a 3T scan generate about 27% higher cortical EF beneath the stimulating electrode than the model based on a corresponding ULF scan. The individual mean relative difference for LMC (radius of 5 mm) shows an error of up to 57% (see Subject 10, **Figure 3A**). For all the 23 subjects, the relative difference is 40% ± 9% (mean ± standard deviation, **Figure 3A**). When considering a larger ROI (radius of 10 mm), the individual mean relative error and the average mean relative errors were ballpark similar (**Figure 3B vs 3A**).

**Figure 2:**
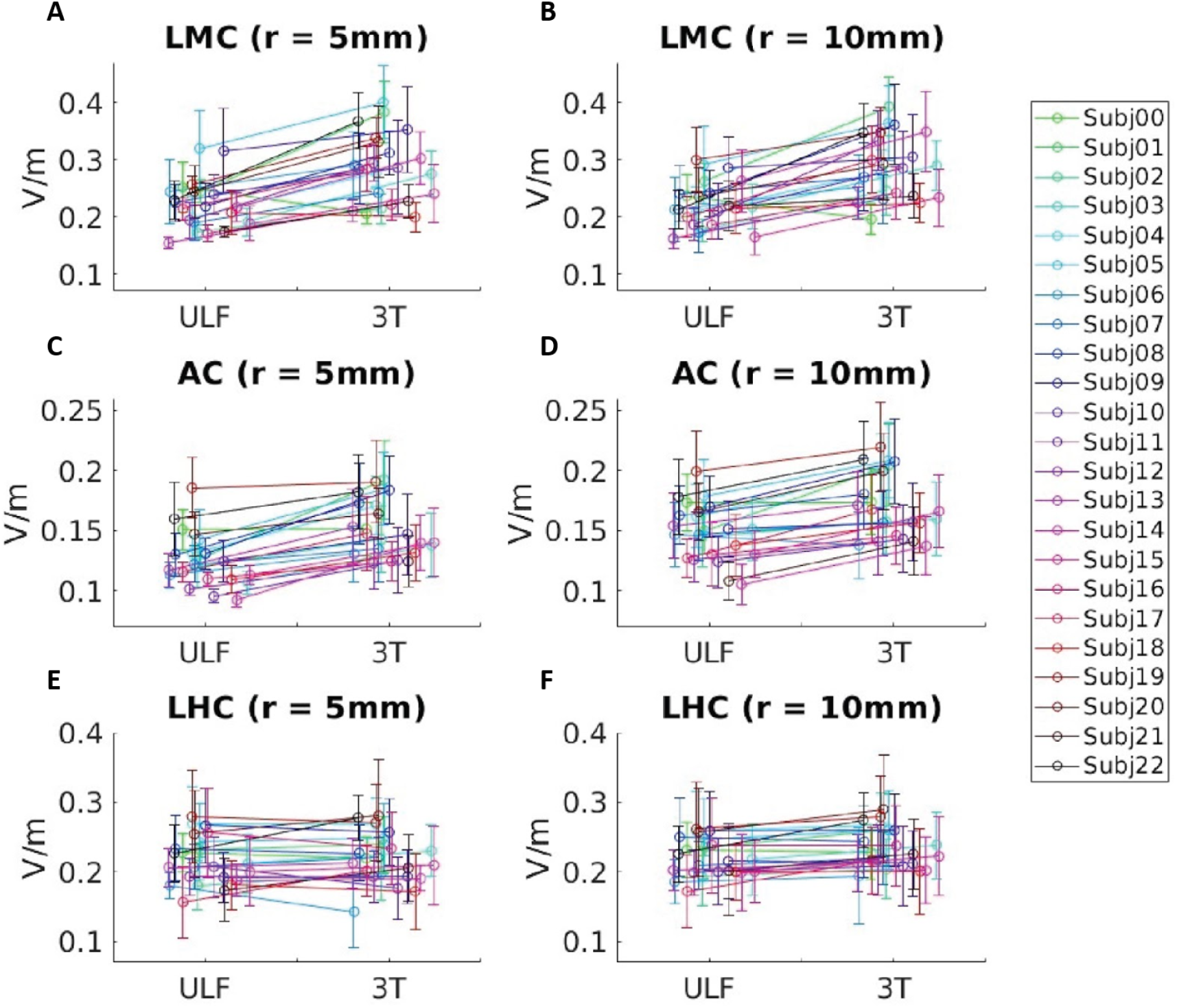
**Induced electric field (EF) magnitude for each of 23 subjects** at the ROIs with different sizes (spheres of 5 mm and 10 mm radii) when considering corresponding ULF and 3T MRI. **Top row:** Left motor cortex (LMC); **Middle row:** anterior commissure (AC); **Bottom row**: left hippocampus (LHC). Each dot is the mean field magnitude in the sphere and the error bar indicates standard deviation. The legend on the right indicates individual subjects.

**Figure 3:**
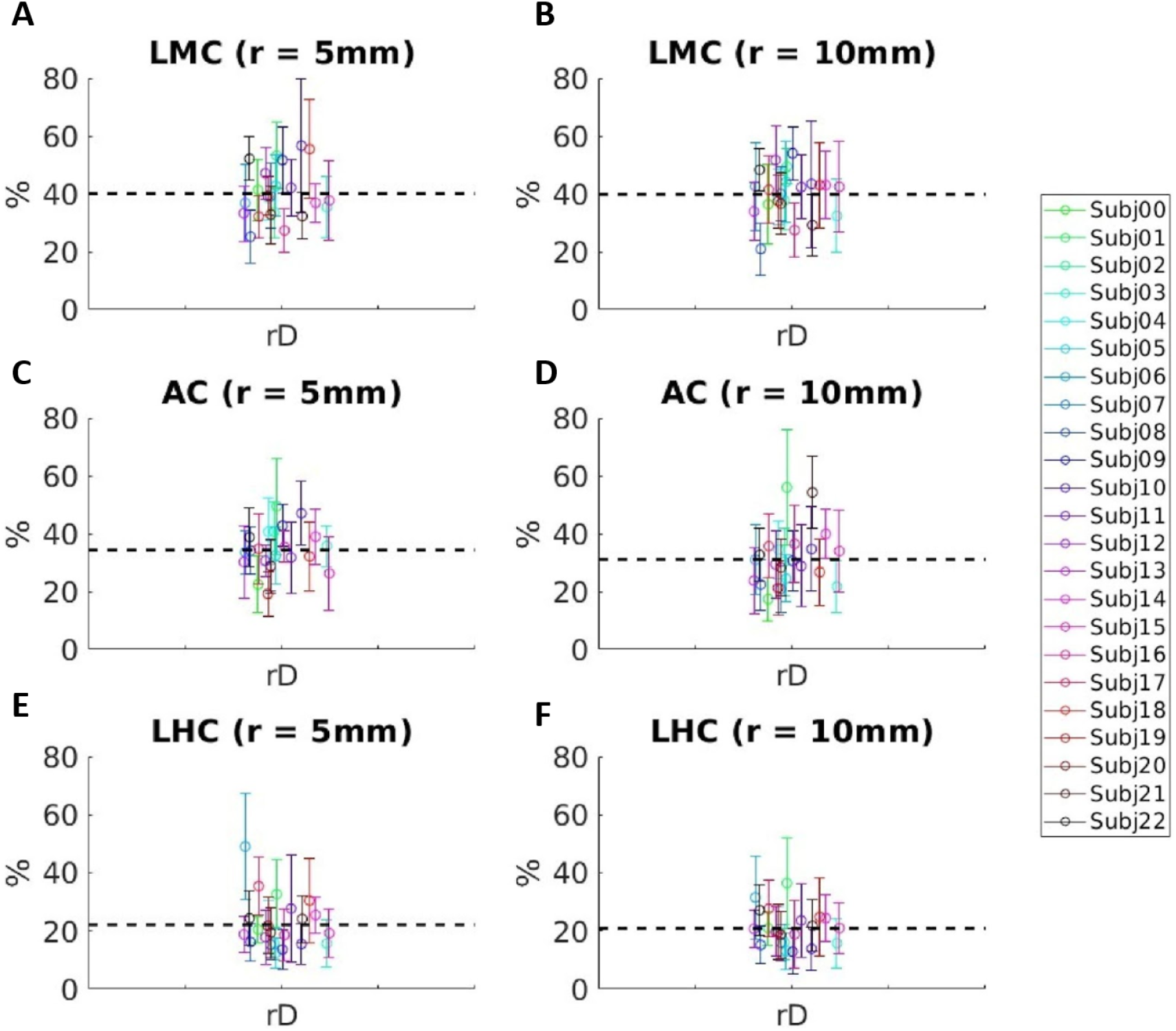
Similar to Figure 2, but showing the relative difference (rD) between the electric field (EF) from ULF and 3T scans. The dashed line at each panel indicates the grand average of rD.

At the ROI of AC, the individual mean EF based on an 3T scan continued to be elevated by about 13% with respect to the corresponding ULF scan (5 mm, ULF vs 3T: 0.12 ± 0.02 V/m vs 0.15 ± 0.02 V/m, **Figure 2C**; 10 mm, ULF vs 3T: 0.15 ± 0.02 V/m vs 0.17 ± 0.03 V/m, **Figure 2D**). However, there is a drop in the mean relative error compared to the first ROI (range: 31–34%) (**Figure 3CD**).

When considering a deeper ROI (LHC), comparison of the individual EF values indicated no clear bias when the ROI size is 5 mm (ULF vs 3T: 0.22 ± 0.03 V/m vs 0.22 ± 0.04 V/m, **Figure 2E**). EF magnitude increases slightly for the most of subjects (19/23) at LHC when considering a bigger ROI (10 mm, ULF vs 3T: 0.22 ± 0.03 V/m vs 0.23 ± 0.03 V/m, **Figure 2F**). However, the vectorial difference is smaller compared to the first two ROIs (mean relative difference of ∼20%, **Figure 3EF**). Further, we note that the spherical volume used to define AC in this study does not encompass the LHC but is superior to it. This therefore suggests an “equalizing effect” where potential differences in EF are balanced out, with increasing brain depth.

## Discussion

In this study, we assessed the suitability of ULF MRI for TES modeling. We essentially compared induced EF due to the classic M1-SO montage in subject-matched datasets across three ROI levels. Our results indicate that there is a substantial difference in predicted EF between an ULF and corresponding 3T scan at the *cortical* level. As conventional tDCS studies (owing to its design) are planned based on targeting a cortical structure, this has important implications. A commonly accepted rule of thumb for the magnitude of induced EF is 0.3-0.5 V/m per mA of injected current-based on computational models (7,20) and intracranial validations (10). Therefore, when considering an ULF scan, researchers may consider “scaling” the induced EF by 27% to match predicted values by a corresponding 3T scan for the same subject. However, the appropriateness of this scale-up would depend on the modeling question being addressed.

The low EF predicted by the ULF scan at the cortical level is likely a combination of two factors. The overestimation of CSF results in current being predominantly shunted away from the underlying cortical structure. Next, our tissue segmentations indicate a greater GM assignment with respect to WM for an ULF scan likely reflecting the reported underestimates of tissue volumes at 64 mT relative to 3T (11). So the ROI for an ULF scan in general consists of more GM in relation to WM in comparison to the same ROI for a 3T scan. This results in a *net* higher conductivity in the ULF ROI in comparison to the same 3T-based ROI. As the induced EF is inversely proportional to tissue conductivity, this explains the lower EF in the ULF scan.

Our results indicate that there is an “equalizing” effect between the ULF and 3T with increasing depth with the brain. The equalizing effect is likely explained by the low presence of CSF and greater tissue correspondence to 3T scan in combination with the increasingly diffuse nature of M1-SO tDCS current-flow patterns at deeper levels. However, if the considered deeper ROI is in immediate proximity to a ventricle, our results may not hold. Due to the error dropping to ∼20% at the level of LHC, we speculate that when the region of interrogation is sufficiently deep, an ULF scan may be appropriate to perform forward modeling only-i.e. for computing EF and visualizing overall current-flow pattern from specified electrode locations and currents. Similarly, we speculate that ULF may have greater suitability for forward modeling in modalities that are focused on deep brain stimulation (21,22). This is in line with the aforementioned greater tissue correspondence noted in morphometric comparison studies with higher dice overlap reported for the GM subcortical and WM regions (∼0.8-0.9) (11).

We note that we did not test the utility of ULF for targeting, i.e. for electrode montage optimization (23). Such approaches are based on maximising intensity or focality at a target ROI. Given the observed differences in induced EF between ULF and 3T scan, one can expect impact but the level of impact is hard to gauge unless studied. We further focused on ROI analysis (24) rather than systematically evaluating differences in current-flow patterns at gyri-sulci levels (7,8). Given the clear overestimation of CSF, we can expect differences. The consideration of ULF scans may have greater implications when considering clinical populations. For instance, if the overestimation in CSF is further compounded in stroke population (25), we can expect greater differences from the corresponding 3T scan in comparison to healthy populations. We further considered one montage that is commonly used clinically. The consideration of a “worst case” montage (e.g. Fpz-Iz montage that results in current traversing a longer path) may have indicated greater differences. Finally, we processed ULF scans as ULF images intrinsically are not compatible/ suitable for brain segmentation tools (25). SynthSR is widely accessible and provides contrast-agnostic image segmentation and performs generative image enhancement. We further note that all morphological brain comparison studies have compared performance of processed ULF scans to 3T.

Current TES administration does not apply an *individualized* scalp intensity with most studies using either 1 or 2 mA. Therefore the mismatch of predicted EF magnitude when using an ULF scan has no consequence for scalp intensity dosing, at this time. Whereas for ECT, consideration of an ULF scan may have a bigger implication for scalp intensity dosing given intensity titration to get to a seizure (26). The lack of individualization of scalp intensity in TES stems both from the fact that: a) target EF magnitude that is “effective” is not defined and b) an accepted method to individualize dose is non-existent. Using individual MRI-guided EF models is one approach to individualize dose, and ULF scans offer a pathway for quicker/wider incorporation due to its favorable cost profile. It is further important to note that some 3T contraindications can be addressed by ULF MRI. For instance, owing to a much lower magnetic field, ULF scanners can be safer for patients with certain types of metallic implants. ULF MRI also typically has lower acoustic noise and more open configurations, which can help alleviate sensitivity to loud sounds, anxiety, and claustrophobia. Next, our results may inform the choice of scanning protocol specific to TES modeling (11). For instance, custom sequences could focus on reducing CSF overestimation for cortical regions of interrogation depending on the question of interest.

In summary, a scale-up factor may be considered when using an ULF scan to estimate cortical EF in some cases. However caution is warranted as each modeling study establishes its own standards for accuracy and precision in accordance with the study objectives. Nonetheless, this initial study is expected to help researchers understand the value and pitfalls of considering ULF scans. Extensive future studies are warranted to test the utility of ULF scans for TES modeling as an alternative to 3T scans.

## Notes

### Competing Interest Statement

The authors have declared no competing interest.

